# Medication and Developmental Stage Shape the Brain-Age Gap in ADHD

**DOI:** 10.64898/2026.01.19.700322

**Authors:** Xiaoyi Zhang, Yuetong Yu, Xiang Liu, Yantong Fang, Jinghua Wang, Sophia Frangou, Yufeng Zang, Ruiyang Ge, Hang Zhang

## Abstract

Identifying robust biomarkers for Attention-Deficit/Hyperactivity Disorder (ADHD) remains challenging. The brain-age gap (BAG), which is the difference between predicted biological brain age and chronological age, has been proposed as a marker of atypical neurodevelopment, yet findings have been inconsistent across studies, limiting its clinical translation. We hypothesized that two major sources of heterogeneity, medication status and developmental stage, contribute to these discrepancies. To test this, we analyzed structural MRI data from a large multi-site cohort (N = 947; ages 7–26 years). Individuals with ADHD were stratified by medication history (medicated vs. unmedicated) and developmental stage (childhood vs. adolescence) and compared with matched typically developing controls. We estimated global- and network-level BAG using publicly available pretrained models. Our results revealed that BAG differences were contingent on medication status, with abnormalities predominantly observed in unmedicated individuals. Developmental stage further moderated the presence of this effect: BAG abnormalities were present significantly in children with ADHD but not in adolescents. Network-level analyses demonstrated spatial specificity, with significant effects in visual, ventral attention, and dorsal attention networks. Together, these findings provide compelling evidence that BAG in ADHD is both medication- and development-dependent, highlighting the need to account for treatment exposure and developmental stage when evaluating BAG as a biomarker and further underscoring the importance of early intervention during childhood.

## Introduction

Attention-Deficit/Hyperactivity Disorder (ADHD) is a common neurodevelopmental disorder (Ayano et al., 2020; American Psychiatric Association, 2013; Polanczyk et al., 2015) affecting approximately 5.9% of children and adolescents (Faraone et al., 2021), characterized by inattention, hyperactivity, and impulsivity (Bush, 2010; Castellanos et al., 2006). Clinical observations indicate that the presentation of ADHD symptoms varies across development; hyperactive-impulsive symptoms typically diminish over time, while inattentive symptoms often persist into adulthood (Biederman, 2000; Kaye et al., 2019; Turgay et al., 2012; Wilens et al., 2009). This developmental variability in symptom expression suggests that the underlying neurobiology of ADHD may involve complex developmental processes.

Neuroimaging research has significantly advanced our understanding of ADHD, consistently demonstrating alterations in brain structure and function relative to typically developing individuals (Albajara Sáenz et al., 2019; Cubillo & Rubia, 2010; Qiu et al., 2011; Rubia et al., 2014; M. Yu et al., 2023). Structural MRI studies, in particular, have identified volumetric alterations in regions such as the prefrontal-striatal circuits, cerebellum, and parietal cortices in individuals with ADHD (Durston et al., 2004; Friedman & Rapoport, 2015; Greven et al., 2015; Hoogman et al., 2017; Lukito et al., 2020; Nakao et al., 2011; Valera et al., 2007). Longitudinal studies further revealed that these alterations are not static but unfold along distinct developmental trajectories (Castellanos, 2002; Chang et al., 2024; Connaughton et al., 2024; Shaw et al., 2007, 2014). While these findings provide a valuable characterization of brain alterations in ADHD, they offer limited insight into how an individual brain deviates from its typical developmental expectations. This motivates approaches that explicitly model typical developmental patterns and yield individualized indices of deviation from normative trajectories.

One such approach is the brain-age gap (BAG) framework. BAG models use machine learning to learn the mapping between neuroanatomical features and chronological age and then estimate an individual’s “brain age” from their MRI data (Cole & Franke, 2017; Franke et al., 2010, 2012). The difference between estimated brain age and chronological age—the BAG—provides a quantitative marker of deviation from normative brain development and maturation (Franke & Gaser, 2019; Gaser et al., 2024; Luders et al., 2016). Although BAG has shown promise across several neuropsychiatric conditions, results in ADHD have been inconsistent, with reports of both accelerated and delayed brain development (Hu et al., 2023; Kaufmann et al., 2019; Kurth et al., 2022; Niu et al., 2022). We propose that two sources of heterogeneity that vary substantially across ADHD cohorts—medication status and developmental stage—are likely to contribute to these discrepant findings, given known treatment effects and pronounced neurodevelopmental changes from childhood through adolescence (Blakemore, 2012; Frodl & Skokauskas, 2012; Lenroot & Giedd, 2006; Nakao et al., 2011; Villemonteix et al., 2015).

Medication exposure is a critical consideration when interpreting BAG findings in ADHD. Psychostimulants—first-line treatments for ADHD—have been associated with normalization of brain structure and function (Nakao et al., 2011; Rubia, Alegria, et al., 2014; Villemonteix et al., 2015), raising the possibility that treatment effects may mask or mimic disorder-related neurodevelopmental differences. Because many prior studies pooled medicated individuals with medication-naive participants, observed BAG differences may reflect an admixture of treatment-related and disorder-specific variations (Cortese et al., 2012; Pereira-Sanchez et al., 2021; Spencer et al., 2013). Developmental stage represents another closely related source of heterogeneity: brain development and maturation follow pronounced non-linear trajectories from childhood through adolescence (Frangou et al., 2022; Shaw et al., 2006; Sowell et al., 2003). Consequently, aggregating individuals across broad age ranges can obscure stage-specific effects that may be present only during particular developmental periods (Blakemore, 2012; Dong et al., 2021; Lenroot & Giedd, 2006; Mills et al., 2016). Together, these design and sampling differences across studies provide a plausible explanation for the variability in reported BAG directions and magnitudes in ADHD literature.

In this study, we used both global and network-level BAG approaches (Y. Yu et al., 2024) to quantify brain development in children and adolescents with ADHD. While most prior BAG studies have focused on global brain BAG measure, estimating BAG within each large-scale functional network may be more sensitive to capture circuit-specific alterations relevant to ADHD and may help localize where developmental deviations are most pronounced (Buckner et al., 2008; Castellanos & Proal, 2012; De La Fuente et al., 2013; Fox et al., 2006). We therefore explicitly tested whether BAG effects vary as a function of medication status and developmental stage. Specifically, we hypothesed that (1) individuals with ADHD would show altered global and/or network-level BAG relative to typically developing controls, and (2) any BAG abnormality would be dependent on both medication status and developmental stage.

## Methods

### Participants

Data for this study were obtained from the ADHD-200 Consortium dataset (http://fcon_1000.projects.nitrc.org/indi/adhd200). The full cohort comprises 973 individuals collected from eight centers: Peking University (PKU), Kennedy Krieger Institute (KKI), NeuroIMAGE Sample (NI), New York University Child Study Center (NYU), Oregon Health and Science University (OHSU), University of Pittsburgh (Pitt), Washington University in St. Louis (WUSTL), and Brown University (BHBU). After excluding 26 individuals from Brown University due to unavailable diagnostic information, the final analytical sample comprised 362 individuals with ADHD [age range: 7.25–20.89 years (mean ± SD: 11.61 ± 4.04); 20.77% female] and 585 healthy individuals [age range: 7.09–26.31 years (mean ± SD: 12.19 ± 3.46); 47.86% female]. Detailed demographic information for all subjects in each dataset is summarized in Supplementary Material Table S1. Each dataset was approved by the research ethics review boards of the respective institutions. Signed informed consent was obtained from all participants or their legal guardians before participation.

### Clinical assessment

In the present study, ADHD symptom severity was assessed using the ADHD Rating Scale IV (ADHD-RS-IV; Demaray et al., 2003; DuPauI, Power, et al., 1998). The ADHD-RS-IV is an 18-item assessment scale adapted directly from the DSM-IV criteria, measuring core ADHD symptoms across three domains: inattention, hyperactivity, and impulsivity. Items are rated on a 4-point Likert scale, yielding a total score (Score of ADHD Index) as well as two subscale scores for inattention (Score of inattentive symptoms) and hyperactivity–impulsivity (Score of hyper/impulsive symptoms), with higher scores indicating greater symptom severity. In addition, information on participants’ medication use was provided, reflecting lifetime use of ADHD-related medication. Both the diagnostic status and medication history were obtained from the ADHD-200 cohort.

### Neuroimaging Acquisition and Preprocessing

Structural T1-weighted MRI scans were acquired at multiple sites using 3T or 1.5T scanners with 3D MPRAGE sequences, with voxel resolution ranging from 0.50 × 0.50 × 1.00 mm³ to 1.30 × 1.00 × 1.33 mm³. Detailed scanning parameters are provided in Supplementary Material Table S2. Images were processed using the standard ‘recon-all’ pipeline of FreeSurfer image analysis suite (version 7.2.0; http://surfer.nmr.mgh.harvard.edu/). Feature extraction was then performed separately for global-level and network-level BAG estimations (see Figure 1a): (1) for global-level BAG, 150 morphometric features were extracted, comprising Desikan–Killiany atlas measures of cortical thickness (n = 68) and cortical surface area (n = 68) (Desikan et al., 2006), regional subcortical volumes (n = 14) based on the Aseg atlas (Fischl, 2004), and the intracranial volume (ICV); (2) for network-level BAG, 2,000 morphometric features comprising Schaefer 1,000-parcel measures of cortical thickness (n = 1,000) and cortical surface area (n = 1,000) (Schaefer et al., 2018) were mapped onto the seven functional networks defined by Yeo et al. (2011).

**Figure 1.**
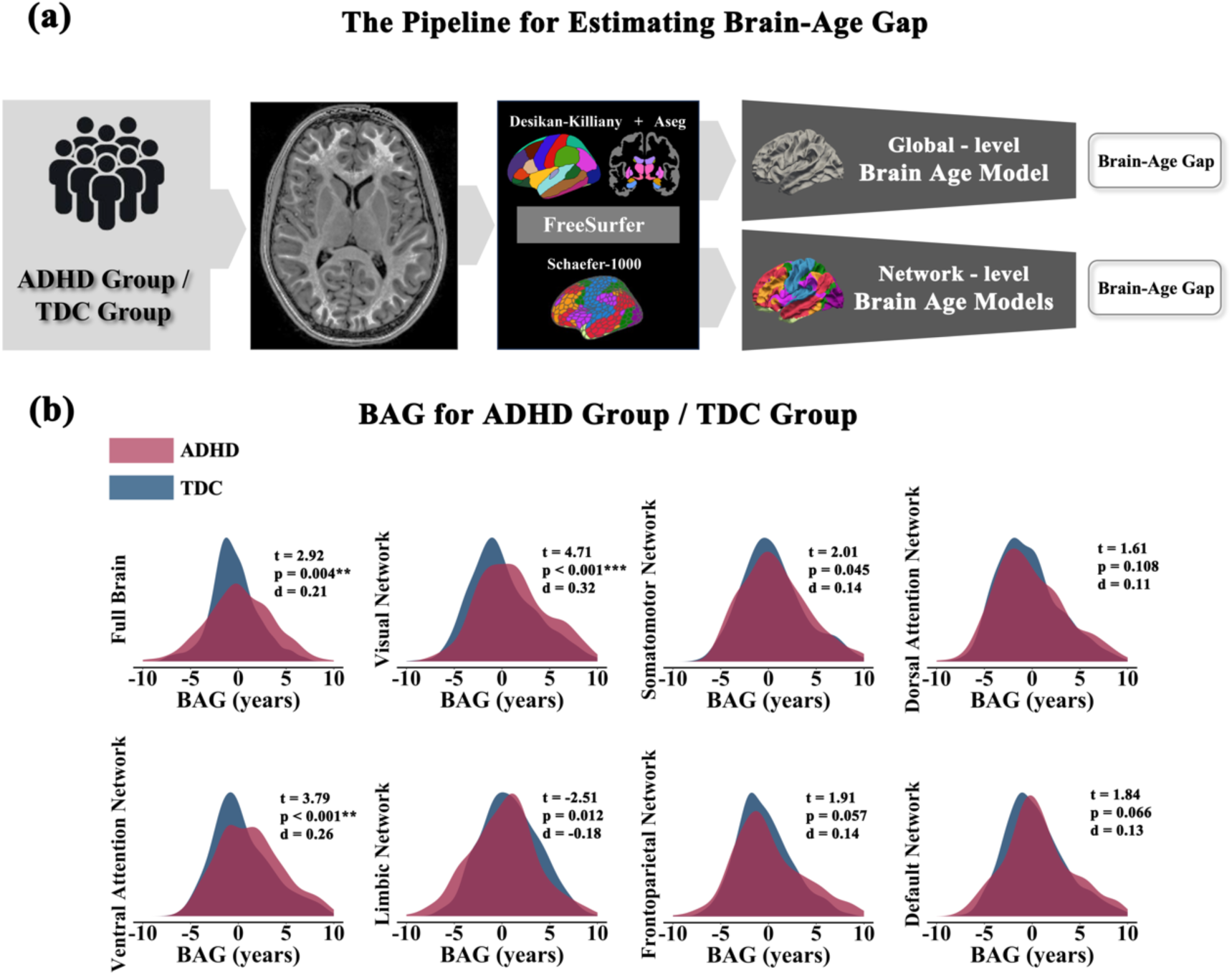
**(a) The pipeline for estimating brain age.** Participants were divided into typical developing controls (TDCs) and individuals with ADHD. Structural MRI scans were processed to generate global-level BAG and network-level BAG. **(b) Group comparisons of global-level and network-level BAG between individuals with ADHD and TDCs.** All results were Bonferroni-corrected for multiple comparisons across networks. **p ≤ 0.01, ***p ≤ 0.001.

### Computation of Global-level BAG

Global-level BAG was estimated using sex-specific pretrained models provided by the CentileBrain framework (https://centilebrain.org/#/) (Ge et al., 2024). These models were developed on FreeSurfer-derived structural features from a large normative sample of 35,683 healthy individuals (53.59% female; age range: 5–90 years) (Y. Yu et al., 2024) and are publicly available via the CentileBrain web platform (https://centilebrain.org/#/brainAge_global). The pretrained model parameters were applied to the current dataset to obtain a global-level BAG estimate for each participant, and to ensure unbiased brain-age values, we used a validated bias adjustment scheme (Beheshti et al., 2019). Positive and negative global-level BAG values indicate apparent older and younger appearing brain, respectively, at the whole-brain level.

### Computation of Network - level BAG

Network-level BAG was also estimated using pretrained models provided by the CentileBrain framework, which are publicly available via the CentileBrain web platform (https://centilebrain.org/#/brainAge_network) (Chakrabarty et al., 2025; Haas et al., 2024). The pretrained model parameters were applied to the current dataset to generate BAG estimates for each of the seven canonical functional networks, including the visual network (VN), somatomotor network (SMN), dorsal attention network (DAN), ventral attention network (VAN), limbic network (LN), frontoparietal network (FPN), and default mode network (DMN). Similar to the global-level BAG estimate, we used a validated bias adjustment scheme to ensure unbiased network-level brain-age values (Beheshti et al., 2019).Positive and negative network-level BAG values indicate apparent older and younger appearing brain, respectively, within each functional network.

### Statistical analyses

Group differences in BAG between ADHD and TDC groups at the global and network levels were tested using independent-samples t-tests, corrected for multiple comparisons (Bonferroni). To examine medication effects, participants with ADHD were categorized as medicated (ADHD-M) or unmedicated (ADHD-UM). One-way ANOVA compared BAG among ADHD-M, ADHD-UM, and TDC groups, followed by post hoc pairwise tests. To examine developmental effects, all participants were divided into childhood (6 ≤ age < 12) and adolescence (12 ≤ age < 18) stages. This 12-year age threshold was chosen as an operational cut-off, informed by prior longitudinal neurodevelopmental research indicating major maturational transitions from late childhood to adolescence (Dong et al., 2021; Lenroot & Giedd, 2006). Group differences (ADHD vs. TDC) in BAG were tested within each stage separately. Within the ADHD group, Pearson correlations assessed associations between BAG and clinical scores (ADHD Index, Inattention, Hyperactivity–Impulsivity). All analyses used Bonferroni correction, with significance set at p < 0.05.

## Results

### Participant characteristics

A total of 947 participants were included in the initial dataset (TDC: n = 585; ADHD: n = 362). One participant was excluded due to missing sex information. Additionally, participants with unsuccessful segmentation or poor image quality were excluded (n = 21 for global-level analyses and n = 5 for network-level analyses). The final samples consisted of 925 participants for global-level analyses (TDC: n = 575; ADHD: n = 350) and 941 participants for network-level analyses (TDC: n = 583; ADHD: n = 358) (see details in Supplementary Figure S1). Within the ADHD group, participants were subdivided by medication status (medicated vs. unmedicated) and by developmental stage (childhood: 6 ≤ age < 12 vs. adolescence: 12 ≤ age < 18). TDC participants were also divided into the same two developmental stages for age-related comparisons. Demographic and clinical characteristics of the ADHD and TDC groups are shown in Table 1, and subgroup characteristics are presented in Supplementary Table S5.

**Table 1.**
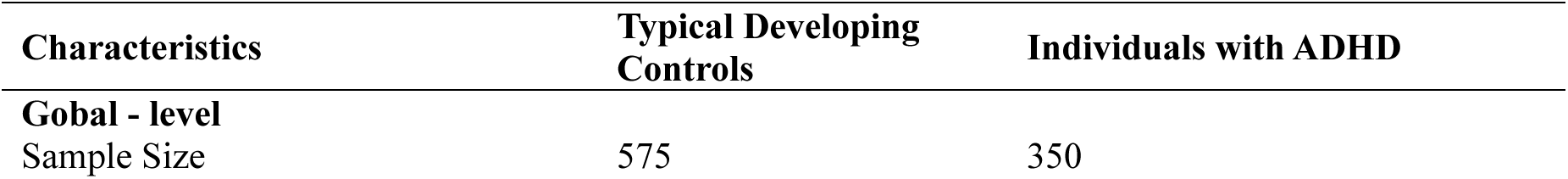
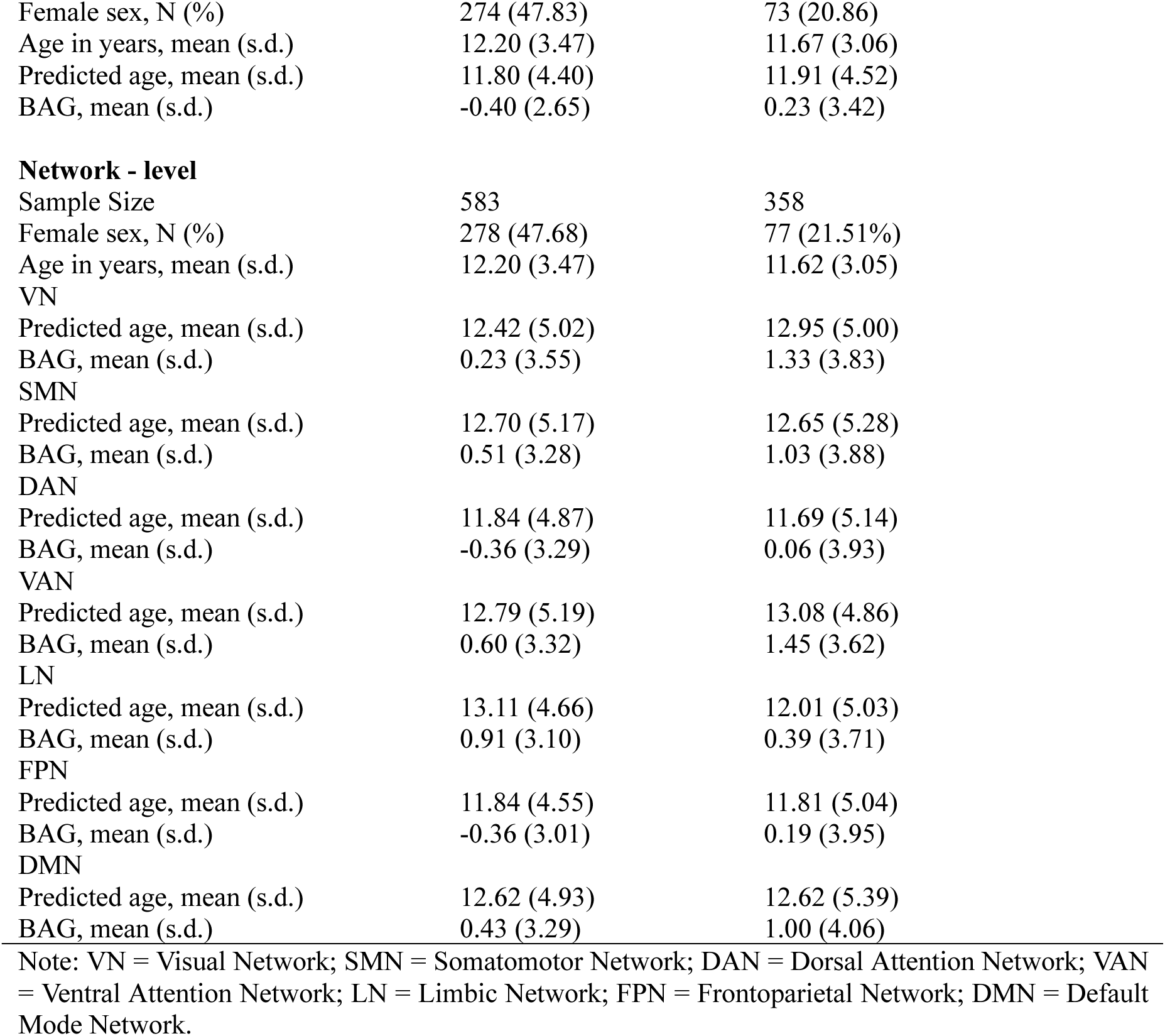
Demographic Characteristics and Brain-age Gap (BAG) Values of Participants by Group.

| Characteristics | Typical Developing Controls | Individuals with ADHD |
| --- | --- | --- |
| **Global - level** |  |  |
| Sample Size | 575 | 350 |
| Female sex, N (%) | 274 (47.83) | 73 (20.86) |
| Age in years, mean (s.d.) | 12.20 (3.47) | 11.67 (3.06) |
| Predicted age, mean (s.d.) | 11.80 (4.40) | 11.91 (4.52) |
| BAG, mean (s.d.) | -0.40 (2.65) | 0.23 (3.42) |
| Network - level |  |  |
| Sample Size | 583 | 358 |
| Female sex, N (%) | 278 (47.68) | 77 (21.51%) |
| Age in years, mean (s.d.) | 12.20 (3.47) | 11.62 (3.05) |
| VN |  |  |
| Predicted age, mean (s.d.) | 12.42 (5.02) | 12.95 (5.00) |
| BAG, mean (s.d.) | 0.23 (3.55) | 1.33 (3.83) |
| SMN |  |  |
| Predicted age, mean (s.d.) | 12.70 (5.17) | 12.65 (5.28) |
| BAG, mean (s.d.) | 0.51 (3.28) | 1.03 (3.88) |
| DAN |  |  |
| Predicted age, mean (s.d.) | 11.84 (4.87) | 11.69 (5.14) |
| BAG, mean (s.d.) | -0.36 (3.29) | 0.06 (3.93) |
| VAN |  |  |
| Predicted age, mean (s.d.) | 12.79 (5.19) | 13.08 (4.86) |
| BAG, mean (s.d.) | 0.60 (3.32) | 1.45 (3.62) |
| LN |  |  |
| Predicted age, mean (s.d.) | 13.11 (4.66) | 12.01 (5.03) |
| BAG, mean (s.d.) | 0.91 (3.10) | 0.39 (3.71) |
| FPN |  |  |
| Predicted age, mean (s.d.) | 11.84 (4.55) | 11.81 (5.04) |
| BAG, mean (s.d.) | -0.36 (3.01) | 0.19 (3.95) |
| DMN |  |  |
| Predicted age, mean (s.d.) | 12.62 (4.93) | 12.62 (5.39) |
| BAG, mean (s.d.) | 0.43 (3.29) | 1.00 (4.06) |

Note: VN = Visual Network; SMN = Somatomotor Network; DAN = Dorsal Attention Network; VAN = Ventral Attention Network; LN = Limbic Network; FPN = Frontoparietal Network; DMN = Default Mode Network.

### Group Differences in BAG between individuals with ADHD and TDC

At the global level, the ADHD group showed significant differences in BAG compared to TDC. Individuals with ADHD exhibited significantly higher global-level BAG compared to TDC (see Figure 1b; *t* = 2.92, *p* = 0.004, Cohen’s *d* = 0.21), indicating older-appearing brain development in the ADHD group.

At the network level, significant group differences were found in the visual network (VN: *t* = 4.71, *p* < 0.001, *d* = 0.32) and the ventral attention network (VAN: *t* = 3.79, *p* < 0.001, *d* = 0.26). No significant differences were found in the other five networks (all *p* > 0.05) (Fig. 1b). All results survived Bonferroni correction.

### Group Differences in BAG by Medication Status

For global-level BAG, a one-way ANOVA revealed a significant group effect (TDC, ADHD-UM, ADHD-M) (*F* _(2, 723)_ = 9.57, *p* < 0.001). Post hoc analyses showed that the ADHD-UM group had a significantly higher BAG than TDCs (*p* = 0.002, *d* = 0.48), whereas the ADHD-M group did not differ significantly from TDCs (*p* > 0.05) (Fig. 2a, Supplementary Table S6).

**Figure 2.**
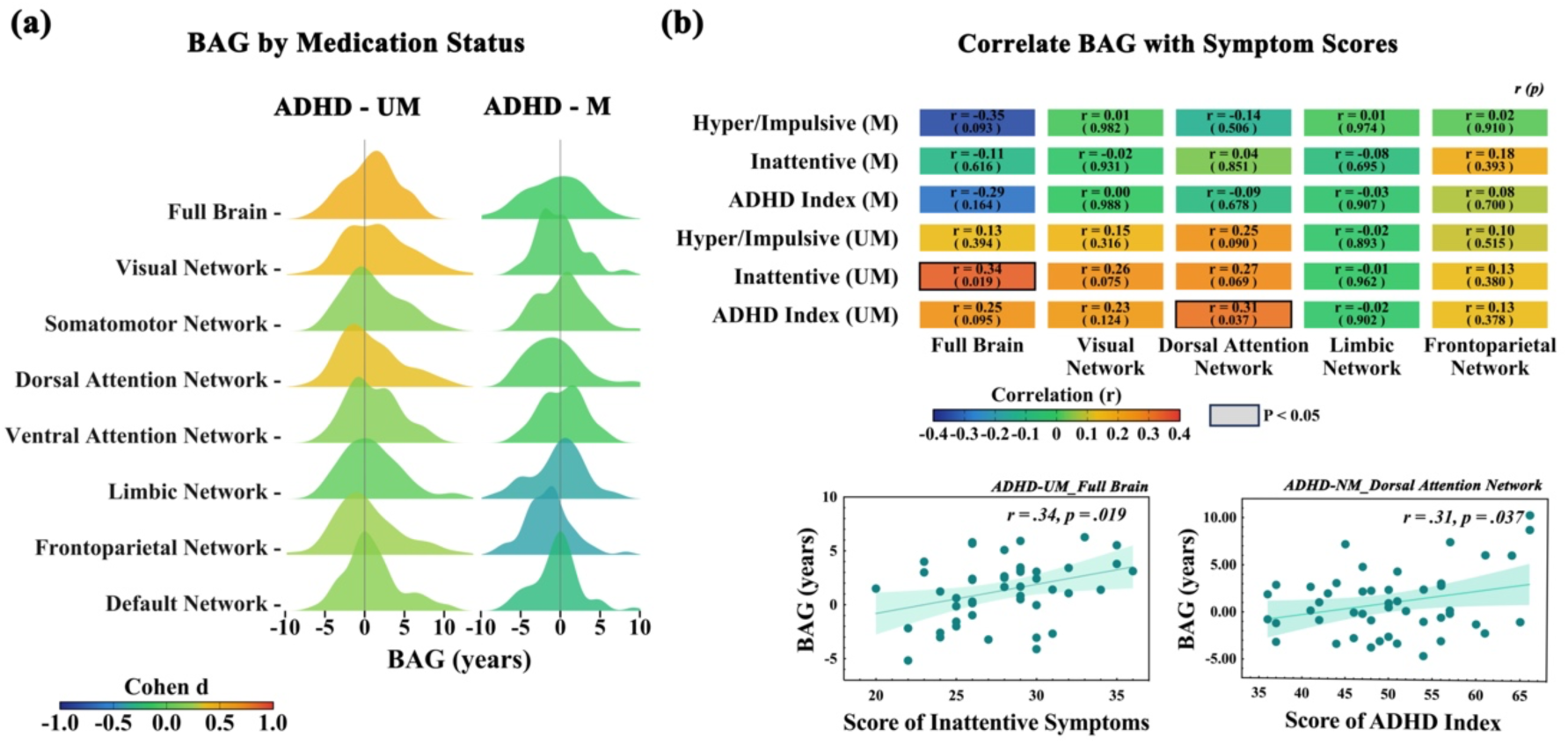
(a) Distribution of global- and network-level BAG for ADHD-UM and ADHD-M participants. (b) Correlations between global- and network-level BAG and clinical symptom scores (ADHD Index, Inattention, Hyperactivity-Impulsivity) in ADHD subgroups. Scatterplots highlighting significant correlations are shown in the lower panels.

At the network level, one-way ANOVAs showed significant group effects in several networks. Post hoc comparisons revealed that, compared to TDCs, the ADHD-UM group exhibited significantly higher BAG in the VN (*p* = 0.001, *d* = 0.41) and DAN (*p* = 0.005, *d* = 0.36). In contrast, the ADHD-M group had significantly lower BAG than TDCs in the LN (*p* = 0.007, *d* = -0.37) and FPN (*p* = 0.005, *d* = -0.38) (Fig. 2a, Supplementary Table S7). All results were Bonferroni-corrected.

Within the ADHD-UM group, global-level BAG was positively correlated with inattentive symptom scores (*r* = 0.34, *p* = 0.019), and BAG in the DAN was positively correlated with ADHD Index scores (*r* = 0.31, *p* = 0.037) (Fig. 2b). No significant correlations were observed between BAG measures and clinical symptom scores in the ADHD-M group (all p > 0.05).

### Group Differences in BAG by Developmental Stage

During childhood, children with ADHD exhibited significantly higher global-level BAG than age-matched TDCs (*t* = 3.86, *p* < 0.001, *d* = 0.37). In adolescence, no significant group difference was observed (*t* = 0.97, *p* = 0.331) (Fig. 3).

**Figure 3.**
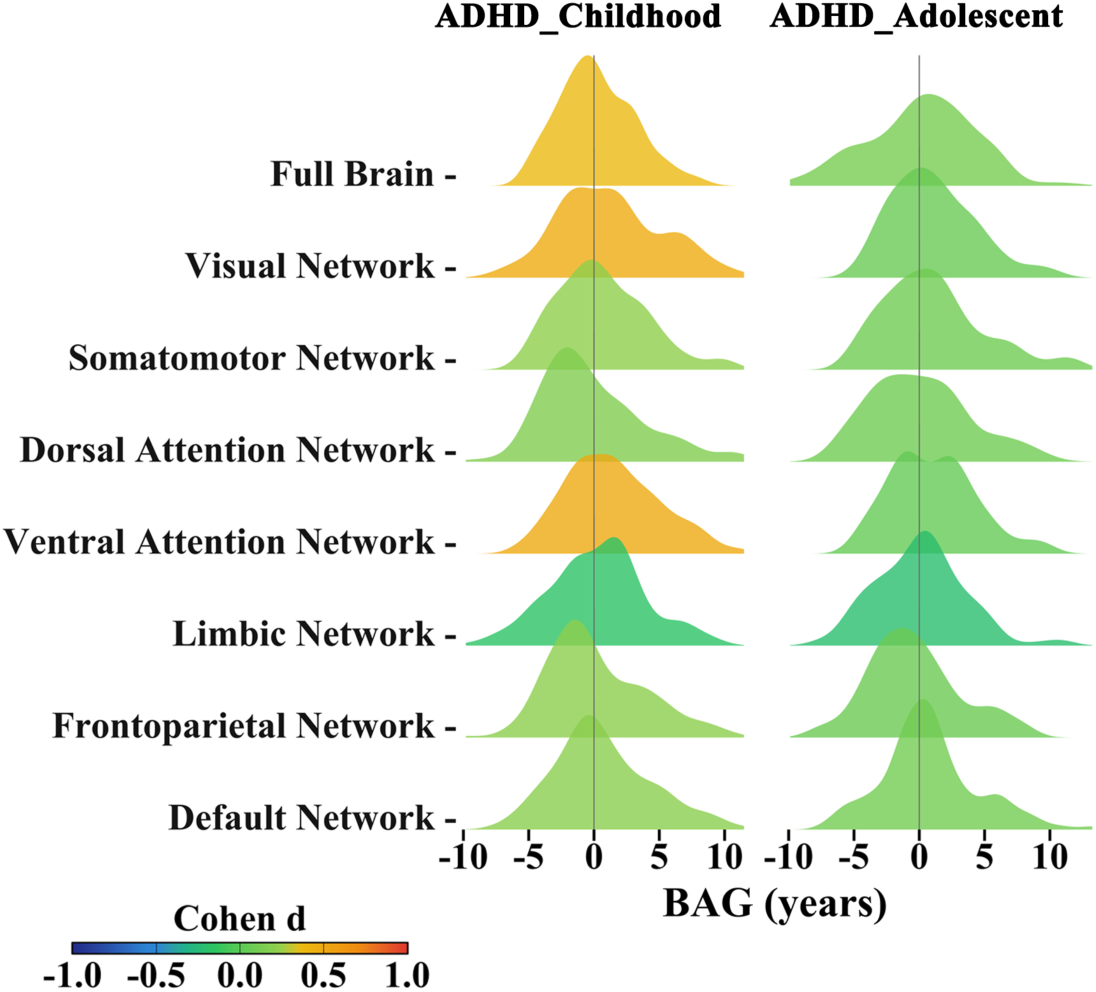
Distribution of global- and network-level BAG for children (6 ≤ age < 12) and adolescents (12 ≤ age < 18) with ADHD. Colors indicate effect size (Cohen’s d) for group differences with age-matched TDC.

At the network level in childhood, significant group differences were observed in the VN (*t* = 5.08, *p* < 0.001, *d* = 0.48) and VAN (*t* = 5.28, *p* < 0.001, *d* = 0.49). No significant differences were found in the other five networks (all *p* > 0.05). In adolescence, no significant network-level group differences were observed between ADHD and TDC groups (all *p* > 0.05) (Fig. 3, Supplementary Table S8). All comparisons were Bonferroni-corrected.

## Discussion

This study utilized a large-scale dataset to investigate brain-age gap (BAG) in individuals with ADHD, focusing on whether BAG abnormalities depend on medication status and differ across developmental stages. Our main findings indicate that significant BAG deviations are present only in unmedicated individuals with ADHD, and only during childhood (6 ≤ age < 12). Second, these abnormalities were not diffuse but showed network specificity, with the largest effects in the visual network, ventral attention network, and dorsal attention network. Collectively, these results suggest that BAG captures a treatment-sensitive deviation from typical brain maturation that is most evident within a time-limited window of early development.

### BAG as a Medication-Sensitive Indicator

The search for biomarkers is often guided by the expectation that candidate markers should track clinical status or treatment response (García De YébenesProus& Carmona Ortells, 2024; García-Gutiérrez et al., 2020; Hussain et al., 2026; Strimbu&Tavel, 2010). In line with this premise, we found no significant BAG deviations in medicated individuals with ADHD: their brain age estimates were statistically comparable to those of typical developing controls. This pattern is consistent with the interpretation that BAG reflects a malleable, treatment-sensitive aspect of ADHD’s neurobiology and could index a normalization process associated with medication. This interpretation aligns with evidence that first-line stimulant medications can induce structural and functional brain changes in children with ADHD (Frodl & Skokauskas, 2012; Nakao et al., 2011; Rubia, Alegria, et al., 2014; Schweren et al., 2013; Spencer et al., 2013; Villemonteix et al., 2015; Wu et al., 2024). More broadly, our findings emphasize BAG as a potential indicator of a medication-modifiable state, underscoring that medication status is a critical design variable for neuroimaging studies of ADHD and should be carefully modeled rather than treated as a nuisance factor or ignored (Cortese et al., 2012; Pereira-Sanchez et al., 2021; Spencer et al., 2013).

### The Developmental Specificity of BAG Deviations

ADHD has long been conceptualized as involving atypical neurodevelopmental trajectories rather than static abnormalities (Dall’Aglio et al., 2022; Noordermeer et al., 2017; Rubia, Alegría, et al., 2014; Shaw et al., 2007). Our finding that BAG abnormalities are confined to childhood and absent in adolescence highlights a critical sensitive developmental window before age 12 —a period marked by rapid and non-linear brain maturation. During this period of brain development (Blakemore, 2012; Lenroot & Giedd, 2006), the brains of individuals with ADHD appear to deviate most conspicuously from typical maturation. This result helps reconcile variability across prior cross-sectional studies spanning wide age ranges: effects that appear inconsistent at the group level may reflect mixing across developmental stages in which deviations are present versus attenuated. The lack of detectable BAG deviation in adolescence could suggest a delayed “catch-up” maturation (Castellanos, 2002; Kakuszi et al., 2020; Linli et al., 2025; Shaw et al., 2007a), compensatory neural plasticity (Larsen & Luna, 2018, 2018; Sharma et al., 2013), or a shift from detectable structural differences to altered developmental trajectories that are less well captured by a cross-sectional BAG snapshot (Castellanos, 2002; Connaughton et al., 2024; Shaw et al., 2007, 2012, 2018). Regardless of mechanism, the restriction of detectable BAG abnormalities to childhood underscores a time-limited neurobiological window, and strengthens the rationale for early intervention, suggesting that therapeutic efforts may be most effective when aligned with this period of peak neurodevelopmental divergence and heightened plasticity (Inguaggiato et al., 2017; Romeo, 2024). This has profound implications, indicating that early childhood may represent a critical period for interventions (Cioni et al., 2016; Hadders-Algra, 2021; Halperin et al., 2012).

### Interpreting the Direction of BAG Deviation: A Composite Measure

A key consideration in interpreting our findings is the nature of the BAG metric. BAG is not a direct measure of the maturity of any single brain region. Instead, it is a composite metric derived from a multivariate model that integrates a wide array of structural neuroimaging features across the brain (e.g., cortical thickness, surface area, volume) (Franke et al., 2010, 2010, 2012; Franke & Gaser, 2019; Kaufmann et al., 2019; Liem et al., 2017). Therefore, a positive BAG (older-appearing brain) should be interpreted as a summary index of distributed, multi-feature structural differences relative to normative aging patterns, not as uniform “acceleration” of development (Whitmore & Beck, 2025). This system-level perspective is supported by our network-level findings, which show heterogeneous network-specific deviations rather than a global shift.

### Network-Specific Alterations

Our network-level analysis revealed that BAG abnormalities were not uniform but were concentrated in visual and attention-related systems, including the ventral and dorsal attention networks. This topography aligns with established evidence implicating these networks in ADHD pathophysiology, particularly for attention regulation and sensory processing (Castellanos & Aoki, 2016; Lin et al., 2021; C. Sripada et al., 2014; Thomson et al., 2022; Tomasi & Volkow, 2012). Notably, the association we observed between network-level BAG and core ADHD symptoms further supports the clinical relevance of the network-level BAG deviations. More generally, the divergent BAG patterns across networks (e.g., medication effects differing by network) are consistent with an imbalanced or dysharmonic developmental trajectory across large-scale brain systems, potentially offering a framework for parsing ADHD heterogeneity beyond whole-brain averages (De Lacy & Calhoun, 2019; Marcos-Vidal et al., 2018; Soman et al., 2023a, 2023b; C. S. Sripada et al., 2014).

### Limitations

Several limitations of this study should be noted. First, while the present results indicate that BAG is sensitive to medication status and developmental stage within ADHD, further work is needed to establish its biomarker properties, including its specificity, sensitivity, and predictive value in independent cohorts. Second, the cross-sectional nature of our data, while identifying a critical window in childhood, cannot delineate the individual-level trajectory of BAG over time. Longitudinal studies are essential to confirm these findings and understand the intra-individual dynamics of brain maturation in ADHD. Third, the imbalanced pattern of BAG across networks suggests complex mechanisms that are not fully captured by regional morphology alone. Integrating functional or structural connectivity measures will be important for testing whether altered inter-network coupling underlies these dysharmonic patterns.

## Conclusion

In summary, our study demonstrates that BAG in ADHD appears to be a dynamic, context-dependent measure: BAG deviations are detectable in unmedicated individuals and are confined to a specific developmental window in childhood. As a composite index of distributed brain structure, BAG provides a complementary systems-level perspective of neurodevelopment in ADHD, particularly when evaluated at both global and network scales. Future longitudinal studies tracking individuals from childhood into adolescence, coupled with multimodal markers (e.g., genetics, cognition, electrophysiology, and brain connectivity), will be critical for clarifying developmental mechanisms and advancing a precision medicine framework for ADHD.

## Author contributions

**Xiaoyi Zhang:** Writing – original draft, Visualization, Software, Conceptualization. **Yuetong Yu:** Software. **Xiang Liu:** Software. **Fangyan Tong:** Funding acquisition. **Jing-Hua Wang:** Funding acquisition. **Sophia Frangou:** Writing – review & editing, Supervision, Funding acquisition. **Yu-Feng Zang:** Writing – review & editing, Supervision, Funding acquisition. **Ruiyang Ge:** Writing – review & editing, Supervision, Software, Project administration, Funding acquisition, Conceptualization. **Hang Zhang:** Writing – review & editing, Supervision, Software, Project administration, Funding acquisition, Conceptualization.

## Acknowledgments

This work was supported by the STI-2030 Major Projects (Grant No. 2021ZD0200500); the Construction Fund of Key Medical Disciplines of Hangzhou, Department of Neurology, Affiliated Hospital of Hangzhou Normal University (Grant No. 2025HZGF02). Jinghua Wang was supported by the Key Research and Development Program in Agriculture and Social Development of Hangzhou Science and Technology Bureau (Grant No. 20241203A12) and the Biomedical and Health Industry Development Support Program of Hangzhou (Grant No. 2022WJC198).

## Data, Materials, and Software Availability

The ADHD-200 data are publicly accessible at [http://fcon_1000.projects.nitrc.org/indi/adhd200]. Models used for the brain-age prediction are available at https://centilebrain.org/#/brainAge_global and https://centilebrain.org/#/brainAge_network.

