## Supplementary Table S1-S8; Figure S1 for "Medication and Developmental Stage Shape the Brain-Age Gap in ADHD"

### S1 Supplementary Methods

**S1.1 Participants**

**Table S1. Demographic information of each dataset in current study.** ADHD = attention-deficit hyperactivity disorder; TDCs = Typical developing controls; KKI: Kennedy Krieger Institute, NI: NeuroIMAGE sample, NYU: New York University Child Study Center, OHSU: Oregon Health Sciences University, PKU: Peking University, Pitt: University of Pittsburgh, WUSTL: Washington University at Saint Louis. SD = standard deviation. More detailed demographic characteristics of participants across sites can be seen in http://fcon_1000.projects.nitrc. org/indi/adhd200.

| Center | ADHD | | | TDCs | | |
| --- | --- | --- | --- | --- | --- | --- |
|  | N | Age Range (Mean±SD) | Gender (F / M) | N | Age Range (Mean±SD) | Gender (F / M) |
| KKI | 25 | 8.10 - 12.99 (10.04 ± 1.55) | 10 / 15 | 69 | 8.02 - 12.97 (10.28 ± 1.26) | 28 / 41 |
| NI | 36 | 11.05 - 20.89 (16.77 ± 2.67) | 5 / 31 | 37 | 11.66 - 20.55 (18.47 ± 3.19) | 25 / 12 |
| NYU* | 151 | 7.24 - 17.61 (10.97 ± 2.67) | 36 / 115 | 111 | 7.17 - 17.96 (12.11 ± 3.10) | 56 / 55 |
| OHSU | 43 | 7.42 - 11.83 (8.96 ± 1.67) | 13 / 30 | 70 | 7.17 - 12.50 (9.20 ± 1.31) | 40 / 30 |
| PKU | 102 | 8.33 - 17.33 (12.09 ± 2.04) | 10 / 92 | 143 | 8.08 - 15.17 (11.42 ± 1.86) | 59 / 84 |
| Pitt | 4 | 13.72 - 17.03 (15.42 ± 1.40) | 1 / 3 | 94 | 10.11 - 20.45 (15.07 ± 2.83) | 44 / 50 |
| WUSTL | 0 | 8.10 - 12.99 (10.04 ± 1.55) | 0 / 0 | 61 | 7.09 - 21.83 (11.67 ± 3.88) | 28 / 33 |
| Totals | 361 | 7.25 - 20.89 (11.61 ± 4.04) | 75 / 286 | 585 | 7.09 - 26.31 (12.19 ± 3.46) | 280 / 305 |

*One subject was excluded due to missing gender information.

**S1.2 Neuroimaging**

**S1.2.1 Acquisition**

**Table S2. Structural MRI acquisition parameters by site.** Seq: imaging sequence, TR: repetition time, TE: echo time, FA: flip angle, TI: inversion recovery delay, PA: parallel acquisition, Res: voxel resolution, SSO: Slice scan order, TA: Total Acquisition Time; BHBU: Bradley Hospital/ Brown University, KKI: Kennedy Krieger Institute, NI: NeuroIMAGE sample, NYU: New York University Child Study Center, OHSU: Oregon Health Sciences University, PKU: Peking University, Pitt: University of Pittsburgh, WUSTL: Washington University at Saint Louis, Trio: Siemens TIM Trio 3T, Allegra: Siemens Allegra, Avanto: Siemens Avanto, MPRAGE: magnetization prepared rapid gradient echo, S: sensitivity encoding (SENSE), G: generalized auto-calibrating partially parallel acquisition (GRAPPA).

| Center | Scanner | Seq | TR (ms) | TE (ms) | FA  (°) | TI (ms) | PA | Res (mm × mm × mm) | SSO | TA |
| --- | --- | --- | --- | --- | --- | --- | --- | --- | --- | --- |
| BHBU | Trio 3T | 3D MPRAGE | 2250 | 2.98 | 9 | 900 | None | 1.00 × 1.00 × 1.00 | Interleaved | 7:36 |
| KKI | Phillips 3T | 3D MPRAGE | 3500 | 3.7 | 8 | 1000 | S ×2 | 1.00 × 1.00 × 1.00 | N/A | 8:08 |
| NIMAGE | Avanto 1.5T | 3D MPRAGE | 2730 | 2.95 | 7 | 1000 | G ×2 | 1.00 × 1.00 × 1.00 | Ascending | 6:21 |
| NYU | Allegra 3T | 3D MPRAGE | 2530 | 3.25 | 7 | 1100 | None | 1.30 × 1.00 × 1.30 | Ascending | 8:07 |
| OHSU | Trio 3T | 3D MPRAGE | 2300 | 3.58 | 10 | 900 | None | 1.00 × 1.00 × 1.10 | Interleaved | 9:14 |
| PKU1 | Trio 3T | 3D MPRAGE | 2530 | 3.39 | 7 | 1100 | None | 1.30 × 1.00 × 1.30 | Interleaved | 8:07 |
| PKU2 | Trio 3T | 3D MPRAGE | 2530 | 3.45 | 7 | 1100 | None | 1.00 × 1.00 × 1.00 | Interleaved | 8:48 |
| PKU3(1) | Trio 3T | 3D MPRAGE | 2000 | 3.67 | 12 | 1100 | None | 0.94 × 0.94 × 1.00 | N/A | N/A |
| PKU3(2) | Trio 3T | 3D MPRAGE | 1950 | 2.6 | 10 | None | 900 | 1.00 × 1.00 × 1.30 | N/A | N/A |
| PKU3(3) | Trio 3T | 3D MPRAGE | 2530 | 3.37 | 7 | None | 1100 | 1.00 × 1.00 × 1.33 | N/A | N/A |
| PKU3(4) | Trio 3T | 3D MPRAGE | 1770 | 3.92 | 12 | None | 1100 | 0.50 × 0.50 × 1.00 | N/A | N/A |
| PKU3(5) | Trio 3T | 3D MPRAGE | 845 | 2.89 | 8 | None | 600 | 1.02 × 1.02 × 1.30 | N/A | N/A |
| PU(3T) | Trio 3T | 3D MPRAGE | 2100 | 3.43 | 8 | None | 1050 | 1.00 × 1.00 × 1.00 | Interleaved | 7:17 |
| WU(3T) | Trio 3T | 3D MPRAGE | 2400 | 3.08 | 8 | G ×2 | 1000 | 1.00 × 1.00 × 1.00 | Ascending | 8:09 |

**S1.2.2 Quality Control**

Using visual inspection, each T1-weighted scan was assessed using four criteria: (1) image affected by movement, (2) temporal poles missing (even partly) in the reconstruction, (3) other parts of the cortex missing in the reconstruction, (4) non-brain tissue (e.g., dural/skull) still visible in the reconstructed pial surface. These criteria were applied separately for the left and right hemispheres. Each criterion was scored 0 if there were ‘no errors visible’ or 1 if there were ‘errors visible in at least three consecutive slices. Scans with scores of 1 or 2 were considered good quality; those with higher scores (>2) were considered of lower quality and were excluded.

**S2 Supplementary Results**

**S2.1 Participant characteristics**

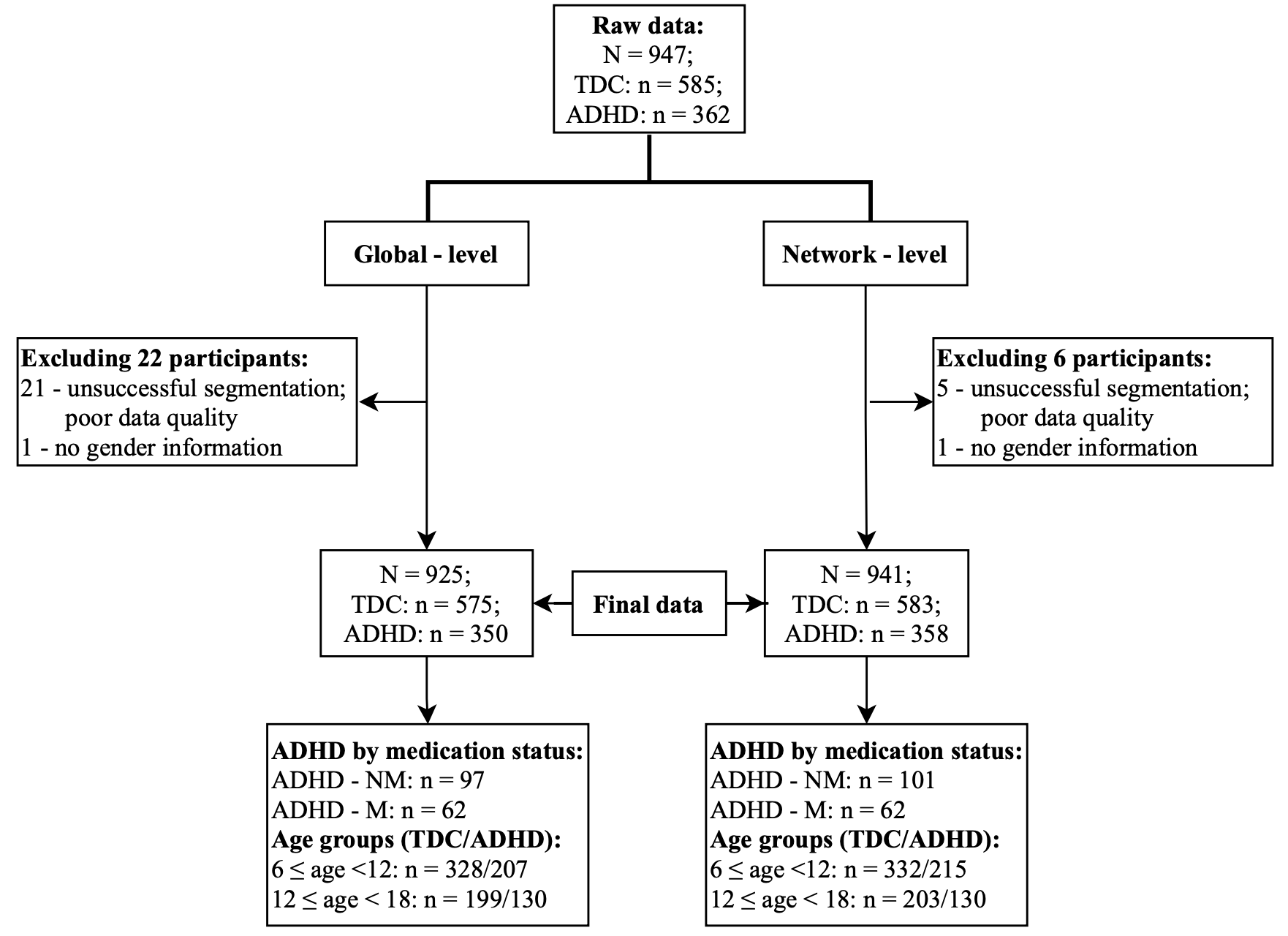

**Figure S1. Flowchart of participant inclusion and exclusion.**

Note: NM = Unmedicated, M =Medicated.

**Table S5 Demographic Characteristics of Participants by Subgroup**

| **Characteristics** | **Typical individuals** | | **Individuals with ADHD** | | | |
| --- | --- | --- | --- | --- | --- | --- |
|  | **Childhood** | **Adolescence** | **Unmedicated** | **Medicated** | **Childhood** | **Adolescence** |
| **Global - level** | | | | | | |
| Sample Size | 328 | 199 | 97 | 62 | 207 | 130 |
| Female sex, N (%) | 158 (48.17) | 86 (43.21) | 21 (21.65) | 9 (14.51) | 54 (26.09) | 17 (13.08) |
| Age in years, mean (s.d.) | 9.78 (1.33) | 14.34 (1.64) | 11.67 (2.37) | 12.24 (2.74) | 9.56 (1.29) | 14.27 (1.60) |
| Predicted age, mean (s.d.) | 9.21 (2.51) | 14.03 (3.14) | 12.61 (4.21) | 11.52 (4.05) | 9.84 (3.15) | 14.42 (4.27) |
| BAG, mean (s.d.) | -0.58 (2.16) | -0.31 (3.19) | -0.94 (3.39) | -0.72 (3.69) | 0.28 (2.96) | 0.15 (4.01) |
| **Network - level** | | | | | | |
| Sample Size | 332 | 203 | 101 | 62 | 215 | 130 |
| Female sex, N (%) | 160 (48.19) | 87 (42.86) | 23 (22.77) | 9 (14.52) | 58 (26.98) | 17 (13.08) |
| Age in years, mean (s.d.) | 9.78 (1.34) | 14.34 (1.65) | 11.60 (2.37) | 12.24 (2.74) | 9.55 (1.29) | 14.27 (1.60) |
| VN |  |  |  |  |  |  |
| Predicted age, mean (s.d.) | 9.60 (3.34) | 15.22 (3.88) | 13.19 (4.57) | 12.25 (4.03) | 10.83 (4.27) | 15.55 (3.83) |
| BAG, mean (s.d.) | -0.17 (3.16) | 0.87 (3.93) | 1.59 (4.08) | 0.01 (3.07) | 1.28 (3.97) | 1.28 (3.52) |
| SMN |  |  |  |  |  |  |
| Predicted age, mean (s.d.) | 9.81 (3.20) | 15.10 (3.72) | 12.74 (4.15) | 12.93 (4.45) | 10.36 (4.01) | 15.49 (4.72) |
| BAG, mean (s.d.) | 0.03 (2.95) | 0.76 (3.54) | 1.15 (3.75) | 0.69 (3.68) | 0.81 (3.67) | 1.22 (4.12) |
| DAN |  |  |  |  |  |  |
| Predicted age, mean (s.d.) | 9.24 (3.38) | 14.11 (3.49) | 12.38 (4.27) | 11.71 (4.40) | 9.50 (4.33) | 14.46 (4.23) |
| BAG, mean (s.d.) | -0.54 (3.09) | -0.23 (3.42) | 0.78 (3.89) | -0.53 (3.61) | -0.06 (4.00) | 0.19 (3.89) |
| VAN |  |  |  |  |  |  |
| Predicted age, mean (s.d.) | 9.87 (3.31) | 15.27 (3.75) | 12.79 (3.74) | 12.67 (4.24) | 11.08 (3.95) | 15.50 (4.13) |
| BAG, mean (s.d.) | 0.10 (3.14) | 0.93 (3.36) | 1.19 (3.54) | 0.44 (3.09) | 1.52 (3.66) | 1.23 (3.47) |
| LN |  |  |  |  |  |  |
| Predicted age, mean (s.d.) | 10.66 (3.21) | 15.09 (3.31) | 12.46 (4.48) | 12.18 (4.47) | 9.87 (4.11) | 14.52 (4.02) |
| BAG, mean (s.d.) | 0.89 (3.08) | 0.74 (3.11) | 0.86 (3.68) | -0.06 (3.60) | 0.32 (3.72) | 0.25 (3.66) |
| FPN |  |  |  |  |  |  |
| Predicted age, mean (s.d.) | 9.43 (2.92) | 13.75 (3.48) | 12.07 (4.58) | 11.02 (4.37) | 9.80 (4.06) | 14.26 (4.51) |
| BAG, mean (s.d.) | -0.35 (2.66) | -0.59 (3.41) | 0.47 (4.25) | -1.22 (3.68) | 0.24 (3.78) | -0.01 (4.16) |
| DMN |  |  |  |  |  |  |
| Predicted age, mean (s.d.) | 9.95 (3.10) | 14.85 (3.83) | 12.82 (4.32) | 12.10 (4.69) | 10.37 (4.19) | 15.32 (4.69) |
| BAG, mean (s.d.) | 0.17 (2.98) | 0.50 (3.62) | 1.22 (3.95) | -0.14 (3.85) | 0.81 (3.90) | 1.05 (4.15) |

Note: VN = Visual Network; SMN = Somatomotor Network; DAN = Dorsal Attention Network; VAN = Ventral Attention Network; LN = Limbic Network; FPN = Frontoparietal Network; DMN = Default Mode Network.

**S2.2 Group Differences in BAG by Medication Status**

**Table S6 Comparison of the global-level BAG by Medication Status**

| Group | N | BAG (Mean ± SD) | Comparison BAG | Post hoc comparisons |
| --- | --- | --- | --- | --- |
| TDC | 575 | -0.40 ± 2.65 | F _[2,723]_ = 9.57 | ADHD-UM > TDC (p=0.002**, d = 0.48) |
| ADHD-UM | 97 | -0.94 ± 3.39 | p<0.001*** | ADHD-M > TDC (p = 0.889, d = -1.12) |
| ADHD-M | 62 | -0.72 ± 3.69 |  | ADHD-UM > ADHD-M (p=0.023, d = 0.45) |

*Significant at p ≤ 0.05 for global-level BAG, or at p ≤ 0.016 for pairwise comparisons among groups

**Table S7 Group Differences in Network-level BAG by Medication Status**

| Network | Groups | BAG | t | p | d | Groups | BAG | t | p | d |
| --- | --- | --- | --- | --- | --- | --- | --- | --- | --- | --- |
|  |  | Mean ± SD |  |  |  |  | Mean ± SD |  |  |  |
| VN | ADHD-UM | 1.59 ± 4.08 | 3.39 | 0.001 | 0.41 | ADHD-M | -0.14 ± 2.84 | -0.63 | 0.531 | -0.08 |
|  | TDC | 0.14 ± 3.40 |  |  |  | TDC | 0.14 ± 3.40 |  |  |  |
| SMN | ADHD-UM | 1.03 ± 3.59 | 1.57 | 0.116 | 0.17 | ADHD-M | 0.43 ± 3.06 | -0.11 | 0.914 | -0.01 |
|  | TDC | 0.47 ± 3.23 |  |  |  | TDC | 0.47 ± 3.23 |  |  |  |
| DAN | ADHD-UM | 0.78 ± 3.89 | 2.88 | 0.005 | 0.36 | ADHD-M | -0.58 ± 3.62 | -0.41 | 0.685 | -0.06 |
|  | TDC | -0.40 ± 3.22 |  |  |  | TDC | -0.40 ± 3.22 |  |  |  |
| VAN | ADHD-UM | 1.05 ± 3.27 | 1.45 | 0.148 | 0.16 | ADHD-M | 0.25 ± 2.76 | -0.67 | 0.502 | -0.09 |
|  | TDC | 0.54 ± 3.24 |  |  |  | TDC | 0.54 ± 3.24 |  |  |  |
| LN | ADHD-UM | 0.86 ± 3.68 | 0.00 | 0.999 | 0.00 | ADHD-M | -0.25 ± 3.30 | -2.71 | 0.007 | -0.37 |
|  | TDC | 0.86 ± 3.02 |  |  |  | TDC | 0.86 ± 3.02 |  |  |  |
| FPN | ADHD-UM | 0.34 ± 4.07 | 1.77 | 0.079 | 0.25 | ADHD-M | -1.51 ± 2.92 | -2.84 | 0.005 | -0.38 |
|  | TDC | -0.41 ± 2.86 |  |  |  | TDC | -0.41 ± 2.86 |  |  |  |
| DMN | ADHD-UM | 0.96 ± 3.53 | 1.72 | 0.087 | 0.19 | ADHD-M | -0.46 ± 2.93 | -1.94 | 0.053 | -0.26 |
|  | TDC | 0.36 ± 3.17 |  |  |  | TDC | 0.36 ± 3.17 |  |  |  |

**S2.2 Group Differences in BAG by Developmental Stages**

**Table S8 Group Differences in BAG by Developmental Stages**

| Network | Childhood _Groups | BAG | t | p | d | Adolescence _Groups | BAG | t | p | d |
| --- | --- | --- | --- | --- | --- | --- | --- | --- | --- | --- |
|  |  | Mean ± SD |  |  |  |  | Mean ± SD |  |  |  |
| VN | ADHD | 1.28 ± 3.97 | 5.08 | 0.000 | 0.48 | ADHD | 1.10 ± 3.23 | 0.56 | 0.573 | 0.06 |
|  | TDC | -0.32 ± 2.89 |  |  |  | TDC | 0.87 ± 3.93 |  |  |  |
| SMN | ADHD | 0.64 ± 3.42 | 2.39 | 0.017 | 0.22 | ADHD | 1.10 ± 3.89 | 0.81 | 0.417 | 0.09 |
|  | TDC | -0.03 ± 2.85 |  |  |  | TDC | 0.76 ± 3.54 |  |  |  |
| DAN | ADHD | -0.12 ± 3.89 | 1.66 | 0.097 | 0.16 | ADHD | 0.09 ± 3.75 | 0.75 | 0.456 | 0.09 |
|  | TDC | -0.64 ± 2.91 |  |  |  | TDC | -0.23 ± 3.42 |  |  |  |
| VAN | ADHD | 1.46 ± 3.55 | 5.28 | 0.000 | 0.49 | ADHD | 1.06 ± 3.24 | 0.36 | 0.720 | 0.04 |
|  | TDC | -0.07 ± 2.85 |  |  |  | TDC | 0.93 ± 3.36 |  |  |  |
| LN | ADHD | 0.27 ± 3.64 | -1.87 | 0.062 | -0.17 | ADHD | 0.07 ± 3.39 | -1.86 | 0.064 | -0.21 |
|  | TDC | 0.83 ± 2.99 |  |  |  | TDC | 0.74 ± 3.11 |  |  |  |
| FPN | ADHD | 0.19 ± 3.70 | 2.18 | 0.030 | 0.21 | ADHD | -0.36 ± 3.53 | 0.62 | 0.534 | 0.07 |
|  | TDC | -0.44 ± 2.49 |  |  |  | TDC | -0.59 ± 3.26 |  |  |  |
| DMN | ADHD | 0.69 ± 3.70 | 2.09 | 0.037 | 0.20 | ADHD | 0.80 ± 3.67 | 0.90 | 0.370 | 0.10 |
|  | TDC | 0.07 ± 2.78 |  |  |  | TDC | 0.44 ± 3.52 |  |  |  |
